# Noise Correlations in Balanced Networks with Unreliable Synapses

**DOI:** 10.1101/2025.01.09.632185

**Authors:** Michael J. Leone, Chengcheng Huang, Brent Doiron

**Author notes:** These authors share senior authorship (Chengcheng Huang, Brent Doiron). (MJL).

## Abstract

Synaptic physiology is highly stochastic in the neocortex: immediately following an action potential, individual synapses release neurotransmitter unreliably, sometimes even failing to release any vesicles. However, theoretical models of neuronal networks typically neglect this well-established feature of biology, especially recurrent networks. In this work, to better understand the effects of synaptic unreliability in recurrent networks, we describe neuronal variability in a balanced network model of non-leaky integrate-and-fire neurons incorporating a Bernoulli model of synaptic release. For arbitrary network size, synaptic unreliability contributes non-negligibly to spike count variability. Most notably, this additional noise is overshadowed by effects on noise correlations. In particular, we find that feedforward and recurrent synaptic reliability have opposite influences on noise correlations: while increased reliability of synaptic input from neurons outside of the network increase correlations, reliability of recurrent synapses de-correlates population activity. We explain this dichotomy by examining the average input currents to cell pairs, and verify this effect with simulations of exponential integrate-and-fire neurons with adaptation and conductance-based synapses. Overall, our results emphasize the importance of synaptic unreliability in the study of noise correlations.

## Introduction

The spike responses of cortical neurons are highly variable across repeated presentations of the same stimulus, such that the trial-to-trial variance of spike counts is comparable to the average number of spikes [1–3]. Studies of neuronal responses *in vitro* have found that neurons fire reliably in response to direct stimulation [4], suggesting that variability predominantly originates from network interaction [5]. In addition, there is a large body of evidence which suggests that this variability is shared across the population, and as a result neuronal pairs exhibit significant noise correlation of their spike trains [6]. With improved techniques for *in vivo* recording of neural populations during behavioral tasks [7], it has been shown that the magnitude of noise correlations is modulated by behavioral state across a wide range of cortical regions as well as species [8, 9, 11]. Both experimental and theoretical work suggest that noise correlations have a significant influence on neural computation [2, 10–12]. However, the origin of variability and noise correlations in the neocortex is still not well understood.

One of the major theoretical advances in the understanding of internally generated variability has been the development of a recurrent model of cortical circuits incorporating the tight balance of strong excitation and inhibition [13–15]. Balanced networks generate chaotic dynamics, reproducing the asynchronous activity typically seen in data of spontaneous firing [15–17]. There is considerable experimental evidence of balance in prefrontal and occipital cortex [18, 19]. Models of dense, balanced networks are characterized by high average spike count variability with spike count correlations that scale inversely proportionally to the number of neurons in the network. The balanced network has served as major advance in the study of intrinsic variability arising in cortical circuits, but other potential sources of variability are well known yet often neglected in modeling.

Spiking statistics reported across cortex may be significantly influenced by a well established, long-known [20] source of synaptic randomness: the unreliable quantity of neurotransmitter release following the onset of an action potential [21–23]. Synaptic release variability has been proposed as a mechanism for rate-dependent spike count variability [24]. Recent work on large-scale networks of detailed biophysical neuron models show that stochastic synaptic transmission contributes predominantly to neural variability compared to other sources of cellular noise [25]. Studies in feedforward pathways have suggested that synaptic unreliability generally de-correlates downstream spike trains [26, 27]. However, there has been little study of how synaptic unreliability affects spike count correlations in recurrent networks. Previous study of recurrently connected purely excitatory [33] or inhibitory [34] networks suggests that synaptic unreliability may cause synchronous firing. The effects of stochastic neurotransmitter release in excitatory-inhibitory networks has not been explored.

In this theoretical study, we first re-examined the overall effects of synaptic unreliability on spike count variability and co-variability in feedforward networks. We found, consistent with previous models [26, 27], that synaptic unreliability always reduces the magnitude of synaptic pairwise correlations. This often leads to an overall input de-correlation of a feedforward input. As a notable exception, we found that if presynaptic excitatory-inhibitory (E-I) pairwise inputs are tightly balanced [15], a decrease in synaptic reliability may actually correlate postsynaptic cell pairs. Next, we considered recurrent networks of non-leaky integrate-and-fire (nLIF) neurons. We showed that with balanced scaling of synaptic strengths, unreliable synaptic release contributes non-negligible variability and co-variability in networks of large population size. Further, we found that while improved reliability of feedforward synaptic release increases the average spike count correlations, improved reliability of recurrent synapses significantly decreased correlations. Last, we demonstrated similar effects of feedforward and recurrent synaptic reliability in networks with more biologically detailed and conductance-based neuron models. These results emphasize the importance of incorporating synaptic unreliability in circuit models in order to understand the mechanistic origin and functional consequences of spiking variability.

## Results

### Vesicle release failure reduces pairwise postsynaptic current correlations

We model vesicle release as an all-or-none process: following an action potential of the presynaptic neuron, each synapse either alters the postsynaptic current of the target cell by a constant amplitude, or fails to elicit a postsynaptic current (Fig. 1A,B). These failures are conditionally independent across synapses given the occurrence of each action potential. Probability of release is treated as a fixed parameter in time and vesicle recovery is instantaneous.

**Fig 1.**
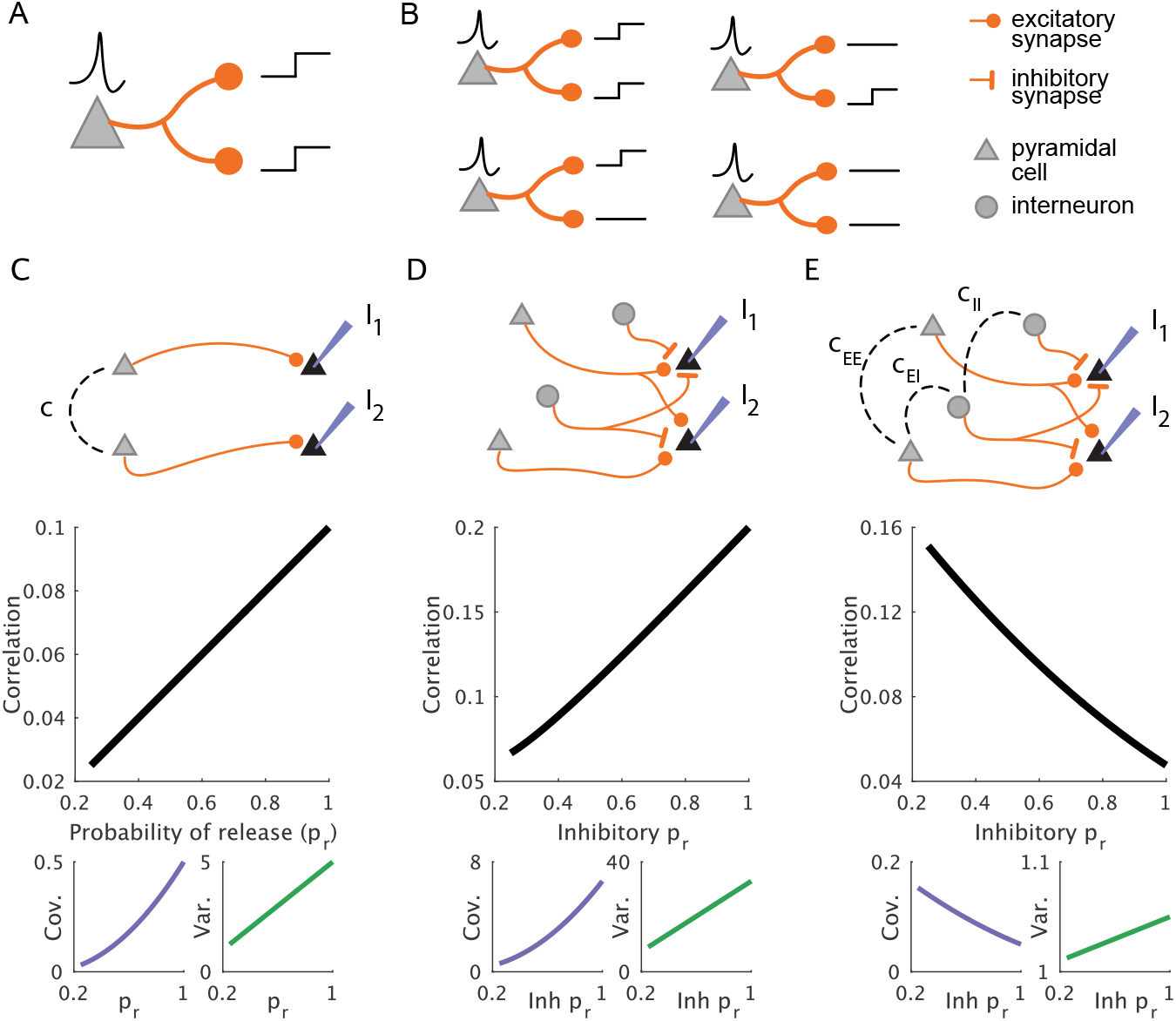
Input correlations due to an external input population with unreliable synapses. (**A-B**) Synaptic failure model description. (**A**) When probability of release is 1, every action potential results in neurotransmitter release at all of that neuron’s synaptic terminals. (**B**) When probability of release is below 1, synaptic terminals act as independent, all-or-none random variables for every action potential. For a given pair of synaptic terminals, there are four alternatives. (**C-E**) Two passive integrators (*I*_1_ and *I*_2_) receive inputs from a presynaptic population of Poisson spiking neurons. (**C**) Input from two correlated neurons. (**Top**) Network schematic. *c* denotes the presynaptic spike count correlation. (**Middle**) Input correlation between *I*_1_ and *I*_2_ as a function of probability of release (*p*_*r*_) of both synapses. (**Bottom**) Covariance and variance plotted separately. (**D**) Input from a population of uncorrelated, densely projecting excitatory and inhibitory neurons.(**Top-Bottom**) Same as in C. (**E**) Input from a correlated population of densely projecting excitatory and inhibitory neurons. (**Top**) Network schematic. *c*_*EE*_, *c*_*II*_, and *c*_*EI*_ denote excitatory-excitatory, inhibitory-inhibitory, and excitatory-inhibitory spike count correlations, respectively. (**Middle-Bottom**) Same as in C and D.

We first consider the simplest case of an individual synaptic pair. If we assume that the length of a trial *T* is much longer than the timescale of a single postsynaptic current, the trial-to-trial variance of the number of successful synaptic transmissions, 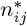, of a single synapse from cell *j* to a target cell *i* satisfies:

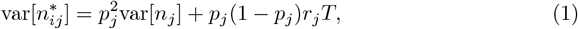

where *p*_*j*_ is the probability of release characteristic of the particular synapse, *r*_*j*_ is the trial-averaged firing rate of neuron *j*, and var[*n*_*j*_] is the variance of the number of action potentials from neuron *j*. The variance of the presynaptic spike count is diluted by a factor 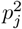 as a direct consequence of thinning the number of effective spikes. In addition to reducing the transfer of pre-synaptic variability, unreliable synaptic transmission introduces a new source of rate-dependent variability, *p*_*j*_(1 − *p*_*j*_)*r*_*j*_*T*. This term corresponds to the variance of the sum of *r*_*j*_*T* independent Bernoulli random variables each with variance *p*_*j*_(1− *p*_*j*_).

For a synaptic pair, there is a similar dilution of output co-variance by reduced probability of release:

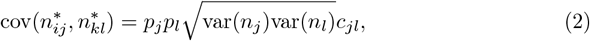

where *c*_*jl*_ is the correlation coefficient of the presynaptic spike counts. This equation describes the case of two correlated but distinct neurons, or of two synapses originating from the same neuron (in which case, *j* = *l* and *c*_*jj*_ = 1). It is apparent that unreliable transmission only reduces pairwise co-variability: random, independent thinning of spikes at each synapse results in a reduction in the number of shared postsynaptic events (Fig. 1C).

By combining these expressions, it is straightforward to show (Eq 22) that the magnitude of the correlation coefficient between synaptic outputs obeys:

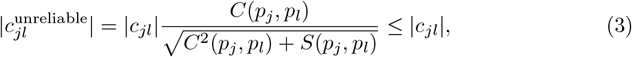

where *C*(*p*_*j*_, *p*_*l*_) and *S*(*p*_*j*_, *p*_*l*_), defined in Eq 22, are non-negative and that *S*(1, 1) = 0, while *S*(*p*_*l*_ < 1, *p*_*l*_ < 1) > 0. From this we see that unreliable vesicle release always reduces the magnitude of pairwise synaptic output correlation compared to the perfectly reliable case.

### Modulation of correlated input depends on the degree of E-I balance

Analysis of the two-synapse case demonstrates that a reduction in synaptic reliability always reduces the magnitude of pairwise input co-variability as well as the correlation coefficient. However, this analysis also applies if the synaptic inputs are anti-correlated. Since neurons receive hundreds or thousands of synaptic connections from interacting excitatory and inhibitory neurons, a given neuronal pair receives a mixture of correlated and anti-correlated inputs. In this section, we demonstrate that changes to input co-variability from a large population of neurons depend critically on the degree of anti-correlation due to excitation-inhibition balance.

We first derive the input statistics to a pair of mutually uncoupled neurons receiving excitatory and inhibitory projections from a large afferent population (Fig. 1D,E, Eq 18 - Eq 28). We simplify the analysis by assuming long-lasting trials, that presynaptic spike counts are homogeneous Poisson processes, and finally by limiting our consideration to an “exact mean field” description such that the number of projections and shared projections to the target cells is the expected value given a dense, random network with connection probability *κ*. We also assume homogeneity of firing rate and synaptic parameters among neurons of either the excitatory or inhibitory type. Each cell receives input from *κN*_*E*_ excitatory and *κN*_*I*_ inhibitory cells, of which there are *κ*^2^*N*_*E*_ and *κ*^2^*N*_*I*_ neurons that comprise a subset of inputs projecting to both cells.

The input variance to a cell is then described as:

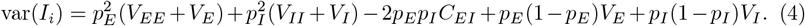

The covariance of the input currents to a cell pair is described as:

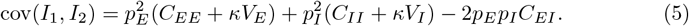

Here, *V*_*E,I,EE,II*_ and *C*_*EE,II,EI*_ (defined in Eq 28) are non-negative coefficients defined in order to encompass all parameters that are independent of probability of release: presynaptic statistics, network density, and synaptic strengths. As two examples, *κV*_*E*_ is the contribution from the proportion *κ* excitatory cells that project to both postsynaptic neurons, and *C*_*EI*_ is proportional to the presynaptic spike count covariance between excitatory cells projecting to one postsynaptic target and inhibitory cells projecting to the other. Since excitatory and inhibitory transmission have opposite effects on postsynaptic currents, positive correlation between E-I spike train pairs causes the term including *C*_*EI*_ to be negative.

In the specific scenario when spike trains of excitatory and inhibitory inputs are uncorrelated, *C*_*EI*_ = 0. Reducing probability of release only leads to a decrease in covariance because all coefficients are positive (Fig. 1D). This is a generalization of the two-synapse analysis (Fig. 1C): all individual synaptic pairs contribute shared events to the postsynaptic currents, and each of these contributions is systematically thinned by decreased reliability. If, however, excitation and inhibition are tightly correlated, *C*_*EI*_ may be large in magnitude (Fig. 1E). With tuned parameters, 2*C*_*EI*_ may be similar in magnitude and opposite to (*C*_*EE*_ + *κV*_*E*_) + (*C*_*II*_ + *κV*_*I*_). By cancellation, total co-variability would be drastically smaller than the individual correlation components, resulting in a reduced average correlation coefficient. This is described in previous work as correlation cancellation [15, 43]. In this case, reduction in probability of release can cause a mismatch of the individual terms and a disruption of the finely tuned relationship between correlation components (Fig. 1E, bottom), leading to a significant increase in average correlation coefficient (Fig. 1E, middle).

### Synaptic unreliability is a non-negligible source of intrinsic variability in balanced networks

In the feedforward framework presented above, we show that synaptic unreliability 1) impacts the transfer of pre-existing variability and co-variability by thinning specific components of input, and 2) acts as a new source of rate-dependent variability due to the randomness of synaptic failure itself (Eq 1). Recent theoretical work proposes the second effect as a mechanism for the high marginal spike count variability widely reported across cortex [24]. In this section, we extend this work by demonstrating that spike count covariances are also impacted by both effects. Particularly in balanced network models, the variability generated by synaptic unreliability itself is non-negligible for arbitrarily large populations. Notably, this is distinct from models where synaptic strength is inversely proportional to the average number of network connections to a neuron. In this case, synaptic noise becomes vanishingly small for large networks, and the only trial-to-trial variability in these networks is imposed by an externally applied noise input.

We consider a recurrent network of non-leaky integrate-and-fire (nLIF) neurons, each receiving independent white noise input with variance σ^2^ (Fig. 2A,B). The covariance matrix of spike counts from time window *T*, as derived in [24], is:

**Fig 2.**
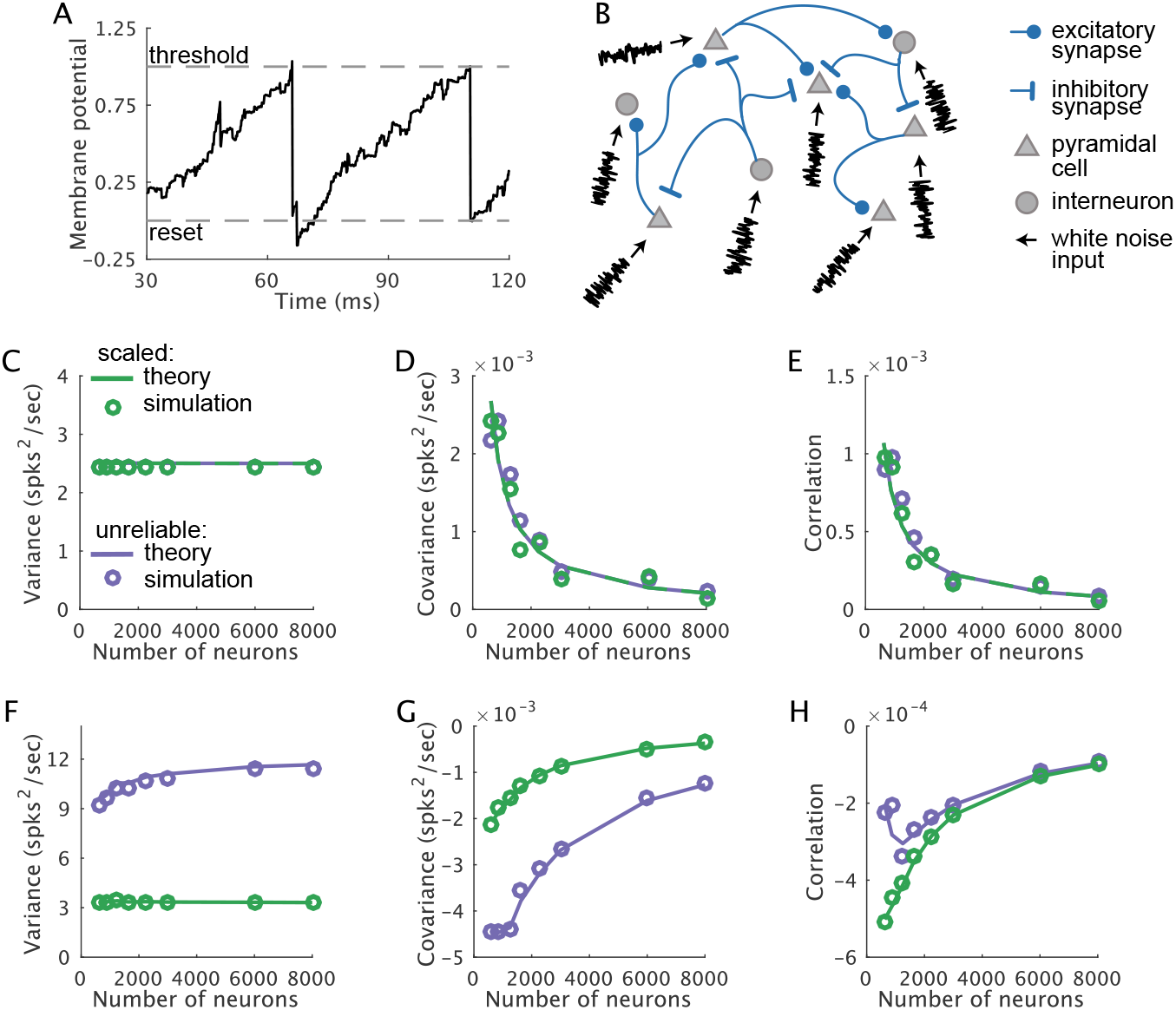
Synaptic unreliability is a non-negligible source of intrinsic variability in balanced networks. (**A**) Sample voltage trace of the non-leaky integrate-and-fire (nLIF) model. When the voltage reaches a hard threshold, there is an instantaneous reset. (**B**) Network schematic. Excitatory and inhibitory neurons are densely coupled and each receives independent white noise. (**C-E**) Networks with 1*/N* scaling, where synapses are perfectly reliable but weakened by half (green, “scaled”), or probability of release is 0.5 (purple, “unreliable”). Theoretical estimation is represented as solid curves and simulation results are in circles. Results are averaged over all cells or cell pairs and over multiple network realizations. (**C**) Spike count variance versus network population size. Spike count covariance versus network population size. (**E**) Spike count correlation coefficient versus network population size. (**F-H**) Same as in (C-E) for balanced networks with 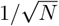 scaling.

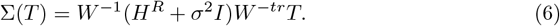

Here *W* is the recurrent coupling matrix where each element, *W*_*ij*_ = *J*_*ij*_*p*_*ij*_, is the trial-averaged synaptic strength from cell *j* to cell *i, I* is the identity matrix, and “ *tr*” is the inverse of the matrix transpose. *H*^*R*^ is the additional variability due to stochastic synaptic release. The diagonal element, 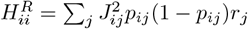, is the sum of the variances of *r*_*j*_ Bernoulli random variables with probability *p*_*j*_ scaled by 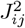 across afferent neurons. The off-diagonal elements of *H*^*R*^ are 0 because synaptic failures are conditionally independent given a set of spikes. We look for network conditions in which *H*^*R*^ is of the same order of magnitude as the external noise variance σ^2^*I*. In this case, changes to probability of release not only affect the transfer of noise through *W*, but also generate significant input variability.

We consider two types of networks with different scaling rules of the synaptic strengths. In the first type of networks, synaptic strength, *J*_*ij*_, scales as 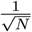, where *N* is the total number of neurons. Networks with such scaling rule, termed as “balanced networks”, generate irregular and asynchronous spiking dynamics [13]. Alternatively, in the the second type of networks, *J*_*ij*_ scales as 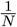, which we refer to as the 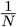 model. We assume both networks are densely connected with fixed probability of connection, *κ*.

We calculate spike count statistics for both network types across a broad range of population sizes. In order to isolate the effects of synaptic randomness on spiking variability, we scale *J*_*ij*_ such that the average synaptic strength, *J*_*ij*_*p*_*ij*_, remains fixed. We performe analysis for two sub-cases: 1) when probability of release (*p*_*r*_) is 0.5, and 2) when synapses are perfectly reliable (*p*_*r*_ = 1) but synaptic strengh are weakened by a factor of 2. Across these sub-cases, firing rates are the same because the trial-averaged input to all cells is invariant. Furthermore, the contribution to variability from externally imposed noise is controlled for since σ^2^*W*^−1^*W*_−*tr*_ is fixed. Therefore, differences in these sub-cases quantify the level of variability generated by synaptic randomness.

We find that in the 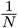 model, across a broad range of population sizes, synaptic scaling and synaptic unreliability have indistinguishable effects on second-order statistics of spike counts (Fig. 2 C-E). Curves of variability, covariability, and correlations of the network with synaptic strength halved are identical to that of the network with synaptic unreliability *p*_*r*_ = 0.5. In contrast, in the network with 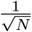 scaling, there are major differences in the network effects of synaptic strength versus synaptic reliability. Population-averaged variance of the unreliable network is more than double that of the network with weakened synaptic strength (Fig. 2F). Similarly, covariances are significantly different in the two networks across a broad range of population sizes (Fig. 2G). Finally, the average correlation coefficients are significantly different for small and moderate population size, and converge in the limit of large population size (Fig. 2H). This suggests that the variance increases in the unreliable networks scale with the increase in the magnitude of covariance.

These results are consistent with the observation that since elements of *H*^*R*^ scale with 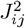, stronger synapses in 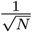 networks should generate larger trial-to-trial fluctuations than those of 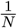 networks. In dense networks, there are on the order of *N* inputs to each cell, such that elements of *H*^*R*^ are proportional to 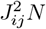, which is invariant to network size in balanced networks with 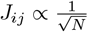.

We make this observation explicit by analyzing an all-to-all network (*κ* = 1) with 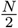 excitatory and 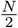 inhibitory cells, and show that only the balanced scaling generates significant input variability. By symmetry, Eq 6 reduces to a pair of scalar equations, each describing the spike count variability of the excitatory or the inhibitory cells (see Supplementary Methods). For brevity, we consider the spike count variability of an excitatory cell. In balanced networks with 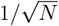 scaling, the synaptic strength between excitatory cells is 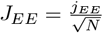, whereas the synaptic strength from inhibitory cells to excitatory cells is 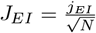, where *j*_*EE*_ and *j*_*EI*_ are independent of network size. For asymptotically large *N*, the variance of the excitatory cell is

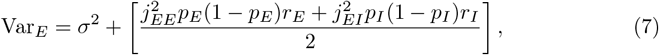

where the second term is the variability due to synaptic unreliability, and *r*_*E*_ and *r*_*I*_ are the firing rates of the excitatory and inhibitory cells. Therefore the variability due to synaptic unreliability remains 𝒪(1) for large population size, meaning that unreliable vesicle release contributes non-negligible variability in dense, balanced networks.

In contrast, in an all-to-all network where 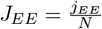 and 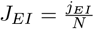, the spike count variance of excitatory cells is

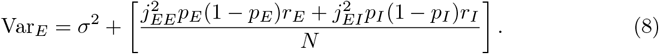

The second term scales as 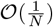, meaning that it vanishes compared to the external noise variance, σ^2^, in large networks.

We see a similar contrast in the two types of networks when considering spike count covariance. In dense, balanced networks, population averaged covariance is inversely proportional to population size. In all-to-all networks, the spike count covariance between excitatory cells is asymptotically

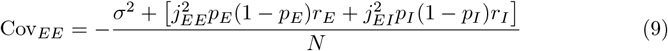

Notably, unlike in the feedforward networks, the added variability due to synaptic unreliability spreads through recurrent coupling, such that the magnitude of co-variability is affected. In the balanced networks, the co-variability arising from external input vanishes at the same rate as the co-variability due to synaptic unreliability. Therefore, synaptic unreliability alters second-order spike count statistics in a way that is distinct from merely changing effective synaptic strength in finite-size balanced networks. However, in networks with of 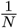 scaling,

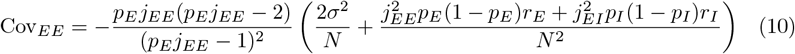

The co-variability due to synaptic unreliability scales as 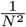, which vanishes much faster than the co-variability due to external noise. The synaptic release probability does modulate the coefficient 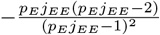 ; however, this is equivalent to weakening synaptic strengths *j*_*EE*_.

In all-to-all networks, the spike count correlation coefficient between excitatory cells is asymptotically

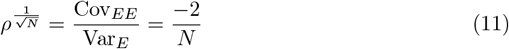

and

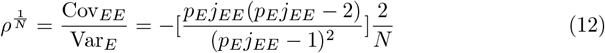

in networks with 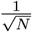 and 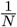 scaling, respectively. Therefore, in large networks, the contribution of synaptic unreliability to the correlation coefficient is only through re-scaling synaptic strength.

The results above suggest that in balanced networks, synaptic unreliability has major effects on variability and co-variability in two separate ways: by changing the effective synaptic strength, and by introducing intrinsic, significant noise. The overall effects on recurrent networks is unclear.

### Unreliability of recurrent synapses increases spike count correlations

We show in Section *Modulation of correlated input depends on the degree of E-I* that probability of release is a major determinant of correlated activity because it affects the degree of input covariance cancellation. In the above Section, we also demonstrated that changes in probability of release have multiple effects in dense, balanced networks. Increased unreliability modifies the average efficacy of synapses, leading to changes in firing rate and the filtering of externally imposed noise throughout the recurrent network. In addition, synaptic unreliability acts as an additional, intrinsic source of input variability. It is therefore unclear how overall correlations are impacted by changes in probability of release for networks of thousands of neurons.

In order to study this, we consider a recurrent network of nLIF neurons that receive random projections from an external population of correlated Poisson neurons (Fig. 3A). This is an extension of the system studied by [24] where we have included correlated external drive. We derive the spike count covariance of neurons over large time windows to be (see derivations in Eq 28 - Eq 34),

**Fig 3.**
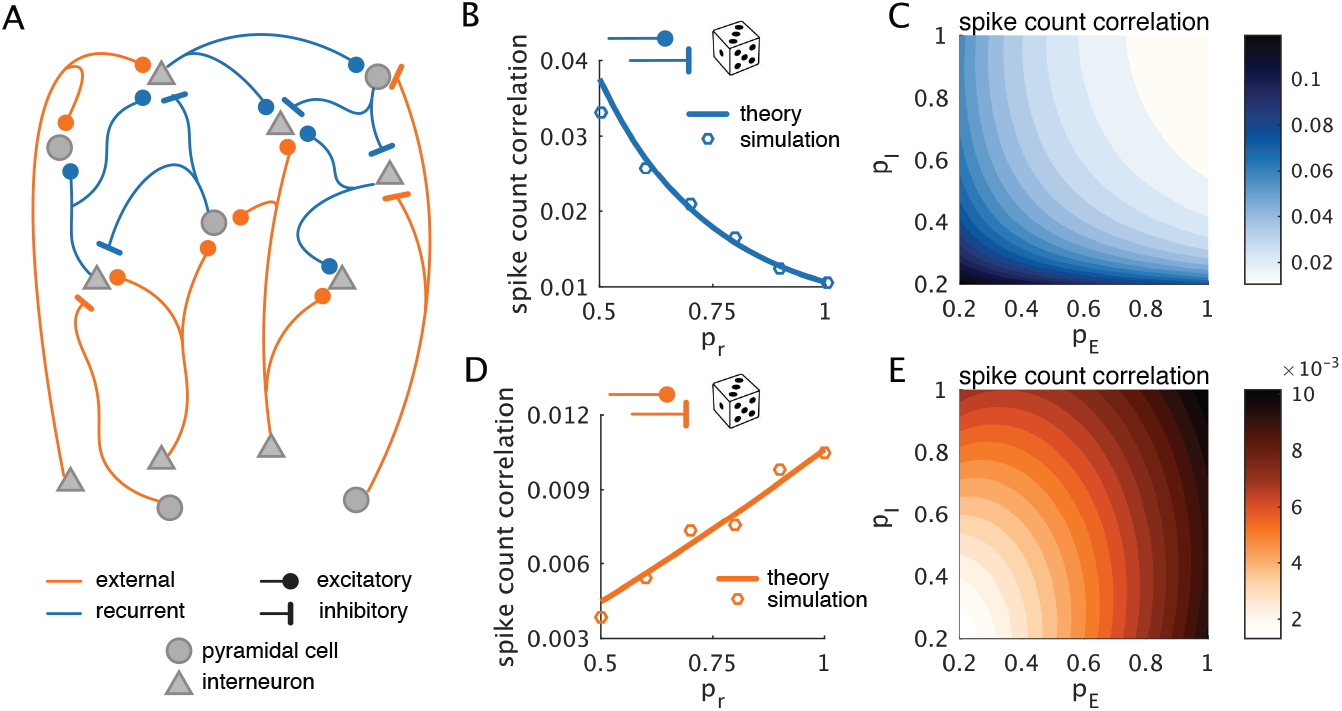
Differential changes in spike count correlations by feedforward and recurrent synaptic reliability. (**A**) Schematic of a recurrently connected spiking network with dense and random projections from an external neuron population. (**B-C**) Changes by recurrent synaptic reliability. (**B**) Population averaged spike count correlations as a function of synaptic probability of release (p_r_) with the excitatory and the inhibitory synaptic release probabilities being equal. Theoretical estimation is represented as solid curves and simulation results are in circles. (**C**) Theoretical estimation of the population averaged spike count correlations for varying excitatory (p_E_) and inhibitory (p_I_) probability of release. (**D-E**) Changes by feedforward synaptic reliability. (**D**) Same as in **B** with p_r_ referring to feedforward synapses. (**E**) Same as in **C** with p_E_ and p_I_ referring to the feedforward excitatory and inhibitory synapses, respectively.

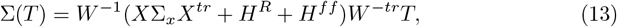

where Σ_*x*_ is the spike count covariance matrix of the external population, a pre-determined parameter of the network. 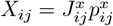 and 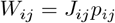 are the feedforward and recurrent coupling matrix, respectively, of trial-averaged synaptic strengths, which are dependent on probability of release. *H*^*R*^ and *H*^*ff*^ represent the additional rate-dependent variability in the network due to recurrent and external neuronal synaptic failures respectively. *H*^*R*^ is defined the same as in Eq 6 as a diagonal matrix with entries 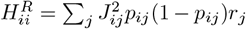. Similarly *H*^*ff*^ is a diagonal matrix with entries 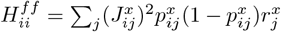, where 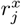 is the firing rate of the external population. We set *T* = 1 without loss of generality. Note that we only consider four synaptic unreliability parameters: *p*_*E*_, *p*_*I*_, 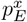, and 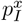, the probability of release of recurrent excitatory, recurrent inhibitory, feedforward excitatory, and feedforward inhibitory synapses, respectively.

We calculate the average correlation coefficient of spike counts in networks with different probabilities of synaptic release and fixed Σ_*x*_ and synaptic weights, *J*_*ij*_ and 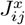. We apply a static bias current to each neuron such that the steady state firing rates are invariant to changes in the probabilities of release. We find that increased probability of release of feedforward synapses results in increased correlations (Fig. 3D,E). In contrast, increased release probability of recurrent synapses results in a large decrease in correlations (Fig. 3B,C). For both types of synaptic reliability, we find that the changes in correlations are predominantly explained by changes in spike count covariance rather than spike count variance (Supplementary Figure S1).

To understand the observed changes in spike count covariance, we consider the various components of input covariance separately. The average input covariance to a pair of cells in the recurrent networks can be broken down as follows (Fig. 4A):

**Fig 4.**
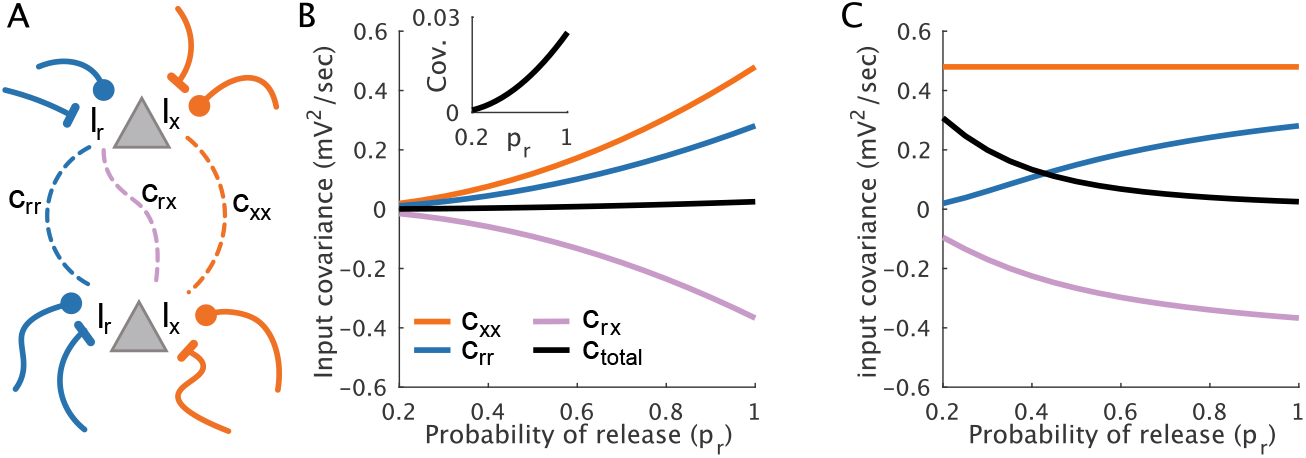
Analysis of input covariance components. (**A**) Schematic of the input covariance components to a pair of cells. Each cell receives input from the external population (altogether denoted *I*_*x*_) as well as from other recurrent neurons (*I*_*r*_). The covariance between the external currents is denoted as *c*_*xx*_, the covariance between recurrent components is *c*_*rr*_, and the covariance between one cell’s external component and the other cell’s recurrent component is *c*_*rx*_. (**B**) The population average of each input covariance component (orange: *c*_*xx*_; blue: *c*_*rr*_; purple: *c*_*rx*_) and the covariance of the total inputs (black), as a function of the probability of release (*p*_*r*_) of feedforward synapses with recurrent synapses being perfectly reliable. Inset: zoomed-in view of the total input covariance. (**C**) Same as **B** for varying *p*_*r*_ of recurrent synapses with feedforward synapses being perfectly reliable.

**Fig 5.**
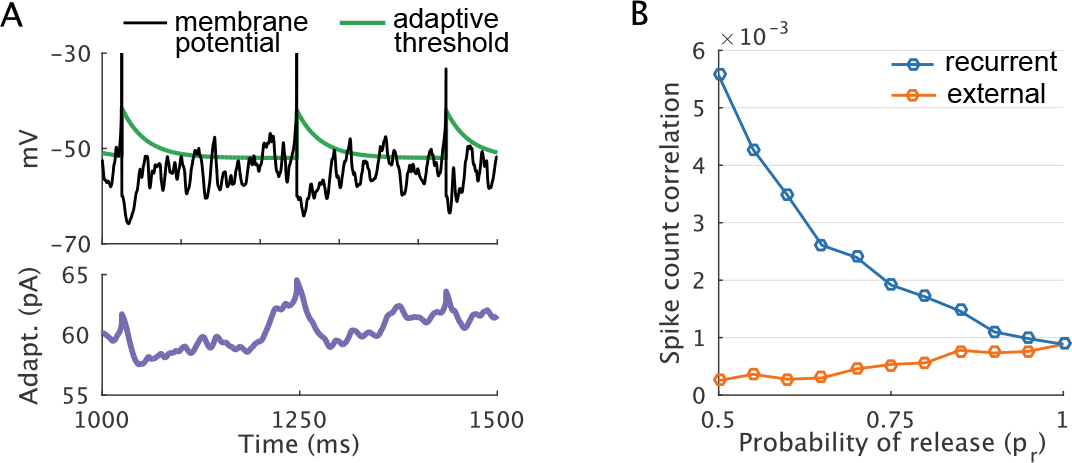
Spike count correlations in a network of conductance-based, exponential integrate-and-fire neurons. (**A**) Sample time traces of a neuron’s membrane potential (black), adaptive spike threshold (green), and adaptation current (purple). (**B**) Population averaged spike count correlation coefficients as a function of the probability of release (*p*_*r*_) of either feedforward (orange) or recurrent (blue) synapses. When probability of release of one type was varied, the other was held at *p*_*r*_ = 1.

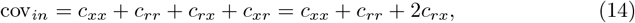

where *c*_*xx*_ is the input covariance due to external currents, *c*_*rr*_ is the input covariance due to recurrent currents, and *c*_*rx*_ is the covariance between recurrent currents to one cell and external currents to another (note that *c*_*rx*_ + *c*_*xr*_ = 2*c*_*rx*_ since we are taking a population average). We expect that feedforward reliability would increase total co-variability by increasing *c*_*xx*_, as is generically the case in the feedforward model (Fig. 1). On the other hand, we expect recurrent synaptic reliability to improve correlation cancellation with increased magnitude of *c*_*rr*_ and *c*_*rx*_, leaving *c*_*xx*_ unaffected.

To confirm this, we derive *c*_*rr*_, *c*_*xx*_, and *c*_*rx*_ analytically (see Methods: *Current Components*) and compare the effects of feedforward and recurrent reliability:

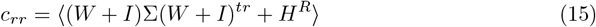

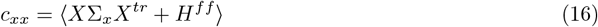

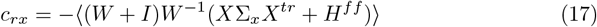

where ⟨.⟩ denotes an average over cell pairs. Indeed, feedforward reliability increases all of the covariance components, leading to an increase in total input covariance (Fig. 4B). In contrast, recurrent reliability improves the efficacy of correlation cancellation through negative feedback, revealed by an increase in *c*_*rr*_ and *c*_*rx*_ (Fig. 4C).

### Correlation modulation via synaptic unreliability in networks of conductance-based, exponential integrate-and-fire neurons

With the nLIF neuron model, we are able to analytically derive spiking and input statistics for arbitrary networks, providing a simplified framework to study the effects of synaptic unreliability on noise correlations. However, there are multiple determinants of noise correlations which arise from intrinsic properties of neurons [8] and unaccounted for in nLIF model, such as nonlinearity of the transfer between input current and response. In addition, synaptic current is simplified as an instantaneous, voltage-independent jump in the previous sections, however, physiologically synaptic current is determined by the membrane potential and membrane conductance, and can last several milliseconds. To confirm that our results apply in a more biologically realistic network, we therefore perform simulations of a dense balanced network with the exponential integrate-and-fire (EIF) spiking neuron model [36]. We also use a conductance-based model of synaptic transmission and incorporate an adaptive spiking threshold and adaptation current, as in[cite Ash]. Finally, there is a 1ms refractory period imposed after each action potential. For the EIF network simulations, we no longer alter the bias current to neurons, and firing rates are free to change with changes in synaptic reliability.

By numerically estimating spike count correlation as a function of either feedforward or recurrent synaptic reliability, again we find that increased reliability of feedforward synapses increases correlations, while recurrent synaptic reliability causes a large decrease. The qualitative consistency between the EIF and nLIF network results suggest a general effect that is expected from a variety of neural models. Because probability of release has such a large effect on input correlation cancellation, other factors such as firing rate and background noise to individual neurons were overshadowed in this particular model.

## Discussion

We have shown that unreliability of neurotransmitter release has differing effects on noise correlations depending on the location and functional role of the synapses. In cases where excitation and inhibition projections are not interacting, synaptic unreliability decreases correlations by independently thinning shared spike signals. Unreliability of synapses which are involved in recurrent feedback will generally increase correlations. With reliable synaptic transmission, large co-fluctuations of excitatory currents can be canceled out with fast inhibitory tracking of excitation [15, 35], while unreliable synapses in recurrent connections unmask the large co-fluctuations in currents and increase spike count correlations. In essence, de-correlation through recurrent feedback is an active, dynamic process that requires strong, reliable interaction, and we find that unreliable synaptic release is detrimental to this process.

In addition to modulating correlations, synaptic unreliability also acts as a source of input variability which is non-negligible in arbitrarily large networks with strong synapses. Contrary to classically scaled networks with synaptic strength inversely proportional to the number of input connections, trial-to-trial synaptic fluctuations in balanced networks have a major, rate-dependent effect on spike count variability. Previous work has shown that unreliable synaptic transmission has the potential to explain large Fano factors across a broad range of firing rates [24]. Our work supplements this theory by isolating the types of models for which this is valid: we found that synaptic variability is significant specifically in the balanced regime.

### Synaptic unreliability is a non-negligible source of trial-to-trial variability particularly in balanced networks

We have shown that the presence of relatively strong synapses with unreliable release dynamics leads to non-negligible, rate-dependent spike count variability in recurrent networks. Therefore, synaptic generation of noise relies on the assumption that postsynaptic potentials constitute a significant percentage of the difference between neuronal resting and threshold potentials. This assumption is supported by both *in vitro* [60, 61] *and in vivo* [62] data, and appears to arise from homeostatic mechanisms [61]. This supports a general theory of internally generated variability largely through intercellular interactions, by way of synaptic unreliability as well as an amplification of external noise through balanced E-I currents. This theory is complementary to other work which shows that clustered architecture contributes additional variability due to slow fluctuations in firing rate [63].

Given evidence suggestive of top-down control of synaptic reliability following stimulus onset [50, 51], it is plausible that increased reliability contributes to the quenching of neural variability following stimulus onset which is observed ubiquitously in different species and cortical regions [64]. However, this intriguing question is beyond the scope of this work.

### Unreliability of recurrent synapses increases spike count correlations

We found that altering probability of release can result in large changes in the correlation coefficient of neural spike trains, reflecting a general property that changes in the functional architecture of a network lead to changes in correlation structure. This phenomenon is separate but not in opposition with theories of correlation modulation in which changes to the marginal statistics of input currents leads to a change in the response gain of neurons [37, 38, 46, 47]. The stochastic nature of neurotransmitter re-uptake, ignored in this work, has also been shown to have a rate-dependent modulatory effect on spike correlations since vesicle recovery is an independent noise source [26]. Our results are instead classified as a modulation of spike count correlation due to a modulation in presynaptic input correlations [8].

Our results suggest a causal relationship between probability of synaptic release and spike count correlations, which is of broad relevance given the ubiquity of state-dependent changes in probability of synaptic release throughout the central nervous system. It is well-known that synaptic depression, a firing rate dependent form of short-term synaptic plasticity, arises due to a reduction in probability of synaptic release [32, 48]. In addition, a recent study in layer 2/3 of rat barrel cortex revealed that GABA_*B*_ receptors [49], presynaptic to pyramidal neurons and activated predominantly by somatostatin (SOM) interneurons, engender a long-lasting modulation of probability of release [50]. Remarkably, noise correlations in rat barrel cortex drop significantly at the onset of whisking [9], concomitant with top-down disinhibition via M1 stimulation of barrel cortex vasoactive polypeptide (VIP) interneurons which in turn inhibit SOM interneurons [51]. Our theoretical results suggest that the increase of pyramidal synaptic probability of release in barrel cortex due to the inhibition of SOM interneurons may contribute to the observed reduction in noise correlations. This phenomenon may also be present in other cortical areas as the VIP-SOM disinhibitory circuit has been found to be a canonical circuit motif across cortex [52–55]. However, further work is needed to better clarify the potential causal relationship between synaptic reliability and behavior-dependent correlation changes.

It is currently unclear why recurrent synapses appear to become more reliable in a state-dependent manner, even though our results suggest that reliability reduces noise correlations, which is generally believed to improve coding performance [2, 10, 11]. It may be that pyramidal cell inactivity and low probability of release during behaviorally inactive epochs preserves energy, given that glutamate release accounts for much of the energy consumption in cortical circuits [56, 57]. More resources are then available to subserve efficient neural processing during appropriate contexts following a top-down signal for behavioral state change. An additional possibility is that high probability of release may not be sustainable over long periods of time due to vesicle depletion.

It is also possible that correlations during spontaneous epochs of activity are beneficial for reinforcement of network structure via long-term potentiation [58]. Increases in probability of release may reduce noise correlations during short epochs opportunistically for coding. Finally, the assumption that a reduction in noise correlation improves coding is only true in particularly special cases, such as in models where neurons have the same tuned response to inputs [12]. Further work is necessary to establish how synaptic reliability influences networks with richer tuning distributions.

In summary, synaptic reliability has a major, and complex impact on variability and correlations at the level of cortical circuits. Our results emphasize the importance of theoretical modeling of synaptic unreliability in the study of intrinsic variability in neuroscience.

## Materials and Methods

### Feedforward network

In this section, we consider the impact of synaptic unreliability on the statistics of the total input to two passive integrators.

Assume that synaptic dynamics are much faster than the total length of a trial, the total current over a period *T* can be treated as a sum of discrete synaptic outputs which are thinned processes of the presynaptic spike trains. The input to a target cell *I* (*i* = 1, 2) is then described as follows:

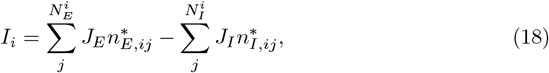

where 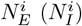 denotes the number of excitatory (inhibitory) cells projecting to cell *i, J*_*E*_ (*J*_*I*_) is the excitatory (inhibitory) synaptic strength, and 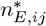 and 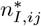 are doubly stochastic random variables, such that given presynaptic spike count 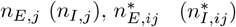 is a binomial random variable with mean 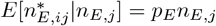 (similarly, 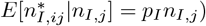. *p*_*E*_ and *p*_*I*_ are the probability of release for excitatory and inhibitory synapses, respectively. Importantly, synaptic outputs conditioned on presynaptic spike counts are independent, meaning that individual synapses do not interact. Spike count correlations between excitatory-excitatory pairs, inhibitory-inhibitory pairs and excitatory-inhibitory pairs are *c*_*EE*_, *c*_*II*_ and *c*_*EI*_, respectively. We assume that spike generation is fixed-rate and population-specific, such that *E*[*n*_*E,j*_] = *λ*_*E*_*T* and *E*[*n*_*I,j*_] = *λ*_*I*_*T*, where *λ*_*E*_ (*λ*_*I*_) is the firing rate of the excitatory (inhibitory) inputs.

The average input to cell *i* is

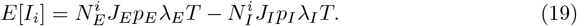

Effective synaptic strength is reduced by synaptic failure in a way that is equivalent to scaling *J*_*E*_ (*J*_*I*_) by *p*_*I*_ (*p*_*E*_).

Since the thinning property of synaptic failure is conditionally independent of presynaptic spike counts, the second order statistics of the input current can be derived such that presynaptic spiking statistics are treated as parameters. The variance of a particular synaptic output 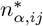 can be broken down as follows:

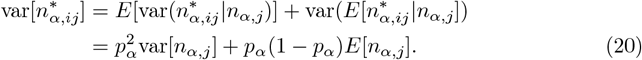

The first term is the dilution of variability by thinning, while the second term is the variance of a binomial distribution with parameters *E*[*n*_*α,j*_] and *p*_*α*_, which accounts for the additional trial-to-trial variability due to the probabilistic nature of synaptic transmission.

The covariance of a particular synapse pair can be expressed similarly:

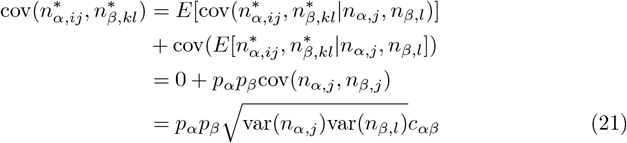

The first term in Eq 21 contributes nothing because synapses are conditionally independent, while the second term is purely a dilution of spike count covariance. This is also true if synapses arise from the same axon, in which case the spike count correlation *c*_*αα*_ is 1. It follows that reduced probability of release always dilutes correlations at the single-pair level:

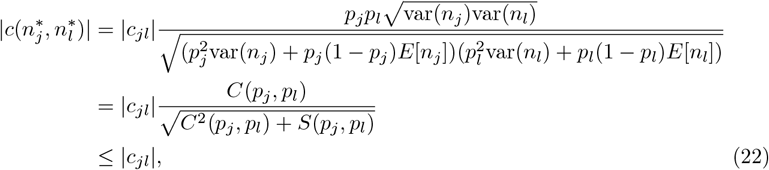

where 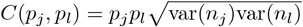. The remainder in the denominator, *S*(*p*_*j*_, *p*_*l*_), is non-negative and positive whenever at least one synapse is imperfect. It follows that the transfer between spike count correlation and input correlation necessarily dilutes as long as at least one synapse is unreliable.

Subsequently, the total variance of the input to cell *i* (from all afferent connections) is calculated directly due to the bi-linearity of variance:

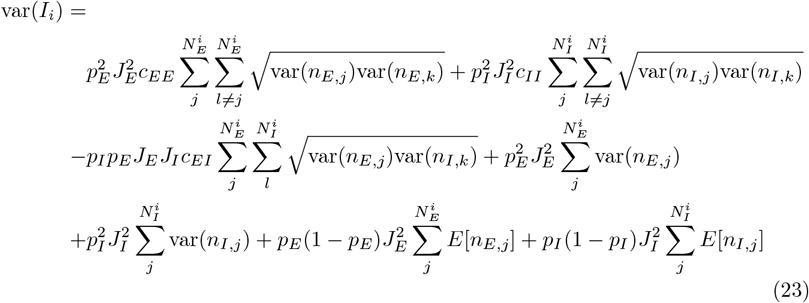

Similarly for the total covariance of inputs to the two cells:

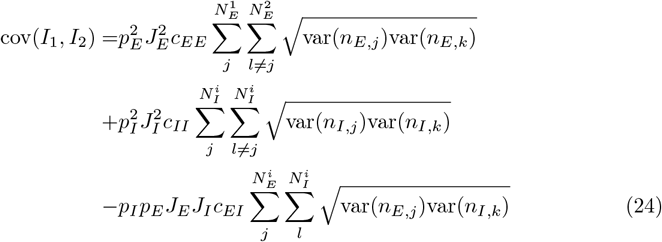

With further assumptions about the connectivity and presynaptic spiking statistics, these expressions simplify considerably. We consider the case when spike trains are homogeneous Poisson processes. In this case, the rate and co-variance are determined fully by spike rate and spike count correlation coefficient. In addition, we employ an exact mean field model where the number of projections and shared projections to the target cells is what is expected on average in a dense, random network with probability of connection *κ*. For a target cell, the expected number of presynaptic excitatory and inhibitory cells is *κN*_*E*_ and *κN*_*I*_, respectively. For a given pair, the expected proportion of shared presynaptic projections is *κ*, so that the number of shared excitatory and inhibitory input neurons is *κ*^2^*N*_*E*_ and *κ*^2^*N*_*I*_, respectively. The expression for variance of synaptic input then becomes

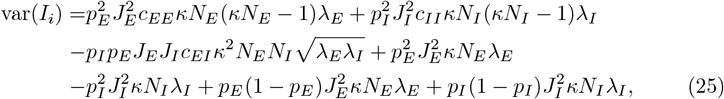

and the covariance between synaptic inputs is

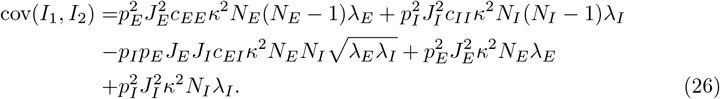

By defining new variables which encompass all parameters besides probability of release, these expressions are greatly simplified:

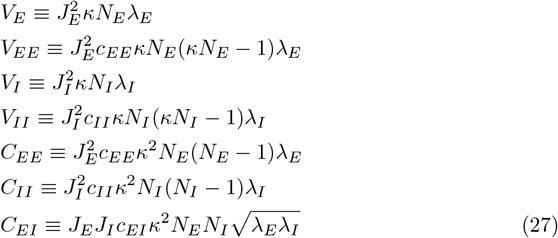

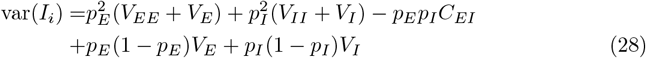

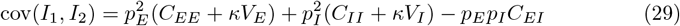

### Recurrent network of nLIF neurons

The spike count covariance matrix of a recurrent network of synaptically unreliable nLIF neurons was previously derived to study the origin of Poisson firing statistics [24], where external input to the network is independent Gaussian white noise applied to each cell. Here we adapt the derivation by replacing the external noise with input from a feedforward network with Poisson firing statistics and random connectivity onto the recurrent network.

The voltage dynamics of an nLIF neuron in the recurrent network is described by the following differential equation:

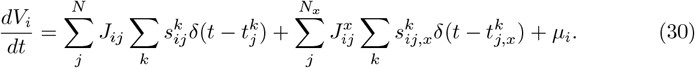

*J*_*ij*_ and 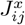 are the coupling strength from cell *j* in the recurrent population or feedforward population, respectively, onto recurrent cell *i*. 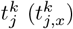 represents the time of the *k*^*th*^ spike from recurrent (feedforward) cell *j*. 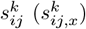 is a Bernoulli random variable with probability *p*_*ij*_ 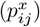, and assumes the value 1 when a synaptic transmission is successful and 0 when the synaptic transmission fails. The firing of the external cells is predetermined to be Poisson processes with rates *r*_*E*_ and *r*_*I*_.

As done in [24], we incorporate the voltage reset rule as self-inhibition. Re-written in vector form,

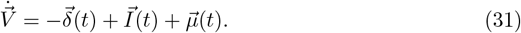

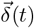 is a vector whose *i*th element is the sum of the *i*th cell’s spike trains, and 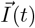 is the total current to cell *i* from other cells in the network. Noting that the average membrane potential is constant in the stationary state and that the synaptic variables are independent of spike times, we take an expectation of Eq 31 over trials and obtain

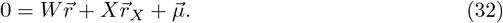

*W* and *X* are random coupling matrices with the same probability of connection *κ*, where 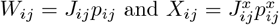. Since elements of 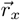 and 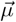 are parameters, one can readily solve for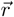. However, in a generic network the input currents can be negative, so the above equation must be modified so that firing rates are non-negative even if input current is negative:

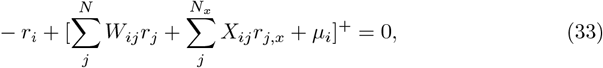

where [*x*]^+^ = *x* if *x* > 0 and 0 otherwise. *r*_*i*_ is therefore the average input current if it is net positive, otherwise *r*_*i*_ is 0. The synaptic unreliability changes the mean firing rates by changing the effective synaptic strengths in *W* and *X*.

Integrating Eq 30 from time 0 to *T* gives

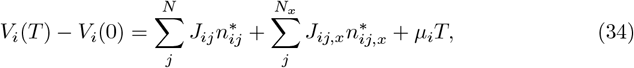

where 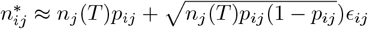and *ϵ*_*ij*_ is an independent normally distributed random variable. Similarly for 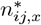. This approximation is accurate for large *T* by the Central Limit Theorem.

Defining Δ*V* (*T*) = *V* (*T*) − *⟨V* (*T*)⟩ and Δ*n*(*T*) = *n*(*T*) − *⟨n*(*T*)⟩, then

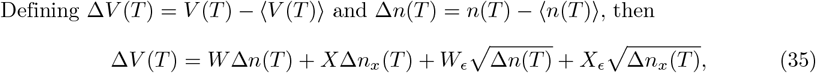

where 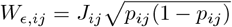 and 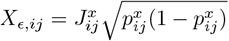. Solving for Δ*n*(*T*)(Δ*n*(*T*))^*tr*^ for large *T* gives

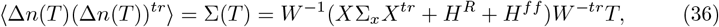

where Σ_*x*_ is the spike count covariance matrix of the feedforward population, which is a pre-determined parameter. *H*^*R*^ and *H*^*ff*^ are diagonal matrices with entries 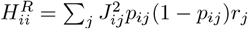 and 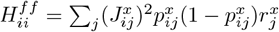, respectively. *H*^*R*^ and *H*^*ff*^ represent additional rate-dependent variability in the network due to recurrent and feedforward neuronal synaptic failures, respectively. Through recurrent interactions, these sources of variability also impact co-variability of spike counts, unlike in the purely feedforward networks.

### Current Components

To better understand how spike count covariance changes with synaptic reliability, we isolate the recurrent and feedforward components of the input current to each cell. The recurrent input fluctuations about the trial average is described as follows:

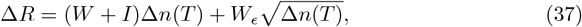

where variables are defined as above. Since the reset rule is not a synaptic current, we have replaced *W* with *W* + *I*. Similarly, the feedforward input fluctuations are

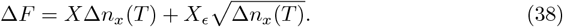

Direct calculation of variance of the recurrent and feedforwad current components yields:

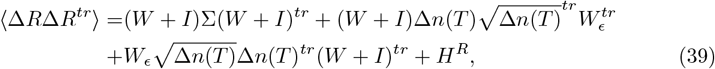

and

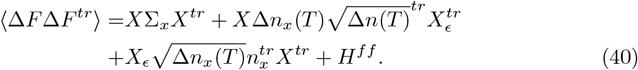

We note that for large *T*, such that Δ*V* (*T*) is negligible, we have the relationship

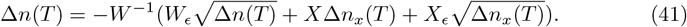

Plugging Δ*n*(*T*) into Eq 39, multiplying by Δ*F*^*tr*^, and taking an expectation, we obtain the unsimplified form of ⟨Δ*R*Δ*F*^*tr*^⟩,

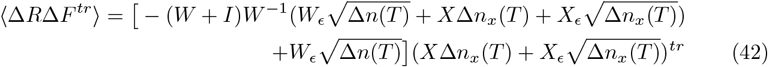

Any terms involving Δ*n*(*T*)Δ*n*(*T*)^*tr*^, 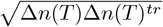, Δ*n*_*x*_(*T*)Δ*n*_*x*_(*T*)^*tr*^, or 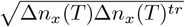 vanish after taking expectations, so covariance terms are as follows:

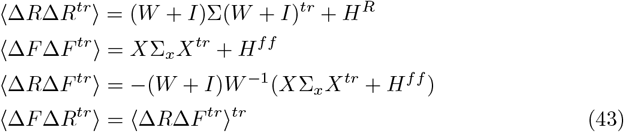

where we have set *T* to 1 second without loss of generality. We identify *c*_*xx*_, *c*_*rr*_, and *c*_*rx*_ as the average of the off-diagonal (cell pair) terms of ⟨Δ*R*Δ*R*^*tr*^⟩, ⟨Δ*F* Δ*F*^*tr*^⟩, and ⟨Δ*R*Δ*F*^*tr*^⟩, respectively.

## Acknowledgments

This work is partially supported by the Intelligence Advanced Research Projects Activity (IARPA) via Department of Interior/ Interior Business Center (DoI/IBC) contract number D16PC00007. The U.S. Government is authorized to reproduce and distribute reprints for Governmental purposes notwithstanding any copyright annotation thereon. Disclaimer: The views and conclusions contained herein are those of the authors and should not be interpreted as necessarily representing the official policies or endorsements, either expressed or implied, of IARPA, DoI/IBC, or the U.S. Government.

## Notes

### Competing Interest Statement

The authors have declared no competing interest.

