## Supplemental Text for "Noise Correlations in Balanced Networks with Unreliable Synapses"

In order to derive the spike count statistics of the all-to-all network, we directly calculated the covariance matrix as in Equation 11. This required inverting the coupling matrix, which has a natural 2x2 block structure:

$$W = \left(\begin{array}{c|c} W_{11} & W_{12} \\ \hline W_{21} & W_{22} \end{array}\right)$$

where $W_{11}$ represents E-to-E connections, $W_{22}$ I-to-I connections, and $W_{12}$ and $W_{21}$ I-to-E and E-to-I connections, respectively.

In general, the inverse of a block matrix exists if and only if $W_{11}$ and $W_{22} - W_{21}W_{11}^{-1}W_{12}$ are themselves invertible. To find the inverse of $W_{11}$, we first express it as the sum of a rank-one and a full-rank (diagonal) matrix:

$$W_{11} = J_{EE}p_E \begin{pmatrix} 1 & \dots & 1 \\ \vdots & \ddots & \vdots \\ 1 & \dots & 1 \end{pmatrix} + \begin{pmatrix} -1 - J_{EE}p_E & \dots & 0 \\ \vdots & \ddots & \vdots \\ 0 & \dots & -1 - J_{EE}p_E \end{pmatrix}$$

Matrices of this form are invertible and there is a known expression for its calculation. We calculate $(W_{22} - W_{21}W_{11}^{-1}W_{12})^{-1}$ using the same method (Sherman–Morrison formula).

The inverse of a full block matrix is expressed as follows:

$$\left(\begin{array}{c|c} W_{11} & W_{12} \\ \hline W_{21} & W_{22} \end{array}\right)^{-1} =$$

$$\left(\begin{array}{c|c} W_{11}^{-1} + W_{11}^{-1}W_{12}(W_{22} - W_{21}W_{11}^{-1}W_{12})^{-1}W_{21}W_{11}^{-1} & -W_{11}^{-1}W_{12}(W_{22} - W_{21}W_{11}^{-1}W_{12})^{-1} \\ \hline -(W_{22} - W_{21}W_{11}^{-1}W_{12})^{-1}W_{21}W_{11}^{-1} & (W_{22} - W_{21}W_{11}^{-1}W_{12})^{-1} \end{array}\right)$$

This is a straight-forward algebraic calculation using the above expressions for $W_{11}^{-1}$ and $(W_{22} - W_{21}W_{11}^{-1}W_{12})^{-1}$. We then express $W^{-1}$ as itself a block matrix:

$$W^{-1} = \left(\begin{array}{c|c} V_{11} & V_{12} \\ \hline V_{21} & V_{22} \end{array}\right)$$

such that $V_{11}$ is $N_E$ x $N_E$, $V_{12}$ is $N_E$ x $N_I$, $V_{21}$ is $N_I$ x $N_E$, and $V_{22}$ is $N_I$ x $N_I$. Recall, the spike count covariance of the nLIF network was derived as follows:

$$\Sigma(T) = W^{-1}(H(r) + \sigma^2 I)W^{-T} \tag{1}$$

Recall also that H(r) is a diagonal matrix where $H_{ii}$ represents the total input variability to cell $i$ due to synaptic unreliability. It is therefore natural to describe H in block structure with the same dimensions as $W^{-1}$.

$$H = \left(\begin{array}{c|c} H_E & 0 \\ \hline 0 & H_I \end{array}\right)$$

$\sigma^2 I$ can also be thought of as a block structure with the same dimensions. By all-to-all symmetry, the input variance to any E cell is the same as to any other (and similarly for I cells) as long as all synapses of the same type have the same unreliability parameter (in this work we only consider $p_E$ ($p_I$) so this is satisfied). As a result, the diagonal elements of $H_E$ ($H_I$) are all the same. Note that input variance to E cells is not the same as to I cells (eg. $J_{EI} \neq J_{EE}$).

$\Sigma(T)$ is therefore calculable as the product of block matrices:

$$\begin{aligned}\Sigma(T) = W^{-1}(H+\sigma^2 I)W^{-T} = &\left(\begin{array}{c|c} H_E V_{11}^2 + H_I V_{12}^2 & H_E V_{11}V_{21} + H_I V_{12}V_{22} \\ \hline H_E V_{11}V_{21} + H_I V_{12}V_{22} & H_E V_{21}^2 + H_I V_{22}^2 \end{array}\right) T \\ &+ \sigma^2 \left(\begin{array}{c|c} V_{11}^2 + V_{12}^2 & V_{11}V_{21} + V_{12}V_{22} \\ \hline V_{11}V_{21} + V_{12}V_{22} & V_{21}^2 + V_{22}^2 \end{array}\right) T\end{aligned} \quad (2)$$

For brevity of exposition, we focus on the spiking statistics of a single cell type, and the most natural choice is excitatory cells and E-E pairs, given that the activity of this cell type is generally expected to be most important for the transfer of information between neural populations. However, qualitative results are similar for other cell types and pairs (not shown). By symmetry, all E cells (and E-E pairs) have the same spike count variance (covariance); it is enough to consider a single diagonal (off-diagonal) element of $H_E V_{11}^2 + H_I V_{12}^2$.

The top left (bottom right) block has two terms, the E (I) variance coefficient and the E-E (I-I) covariance coefficient. "diag" will refer to the variance term and "offdiag" will be for the covariance term. For balance (asymptotically),

$$V_{12}^2(\text{offdiag}) = \frac{j_{EI}^2}{(j_{EE}j_{II} - j_{EI}j_{IE})^2 N^2} + O(\frac{1}{N^{\frac{5}{2}}}) = O(\frac{1}{N^2})$$

which is much smaller than $V_{11}^2$:

$$V_{11}^2(\text{offdiag}) = -\frac{1}{N} + O(\frac{1}{N^{\frac{3}{2}}})$$

Finally, the diagonal term of $V_{12}^2$ is the same as the off-diagonal term which is $O(\frac{1}{N^2})$, and $V_{11}^2(\text{diag}) = 1$, so only the latter matters for large N.

As for the $\frac{1}{N}$ terms,

$$V_{12}^2(\text{offdiag}) = V_{12}^2(\text{diag}) = \frac{j_{EI}^2}{(j_{EE}j_{II} - j_{EI}j_{IE} - j_{EE} - j_{II} + 1)^2 N} + O(\frac{1}{N^2}) = O(\frac{1}{N})$$

$$V_{11}^2(\text{offdiag}) = -\frac{j_{EE}(j_{EE}-2)}{(j_{EE}-1)^2 N} + O(\frac{1}{N^2}) = O(\frac{1}{N})$$

$$V_{11}^2(\text{diag}) = 1 + O(\frac{1}{N})$$

By symmetry, all excitatory neurons have the same firing rate, and similarly all inhibitory neurons have the same firing rate. The subpopulations will not have the same firing rate because of population-specific synaptic weights as well as the difference in external input magnitude. Note the following rate expressions are for asymptotically large N:

$$r_E = \frac{(\mu_I J_{EI} - \mu_E J_{II})}{N p_E (J_{EE} J_{II} - J_{EI} J_{IE})}$$

$$r_I = \frac{(\mu_E J_{IE} - \mu_I J_{EE})}{N p_I (J_{EE} J_{II} - J_{EI} J_{IE})}$$

The diagonal elements of $H_E, H_I$ is in a simple enough form, and we abuse notation again by ignoring that these are matrices and instead refer to the diagonal values:

$$H_E = N_E J_{EE}^2 p_E (1 - p_E) r_E + N_I J_{EI}^2 p_I (1 - p_I) r_I$$

Note that for balanced scaling, $H_E$ is O(1) and for 1/N scaling $H_E$ is O(1/N). For either scaling, the variance of excitatory cells is given by:

$$Var_E = (H_E + \sigma^2) V_{11}^2 (diag) = H_E + \sigma^2$$

Since $\sigma^2$ is fixed, it is clear that for large N, $H_E$ is either negligible for 1/N scaling, or in the balanced case, remains significant for arbitrary N. Unreliability in this sense has no effect on large 1/N scaled networks but is significant in the balanced state. Covariance is affected similarly.
