## Supplemental Figures for "Noise Correlations in Balanced Networks with Unreliable Synapses"

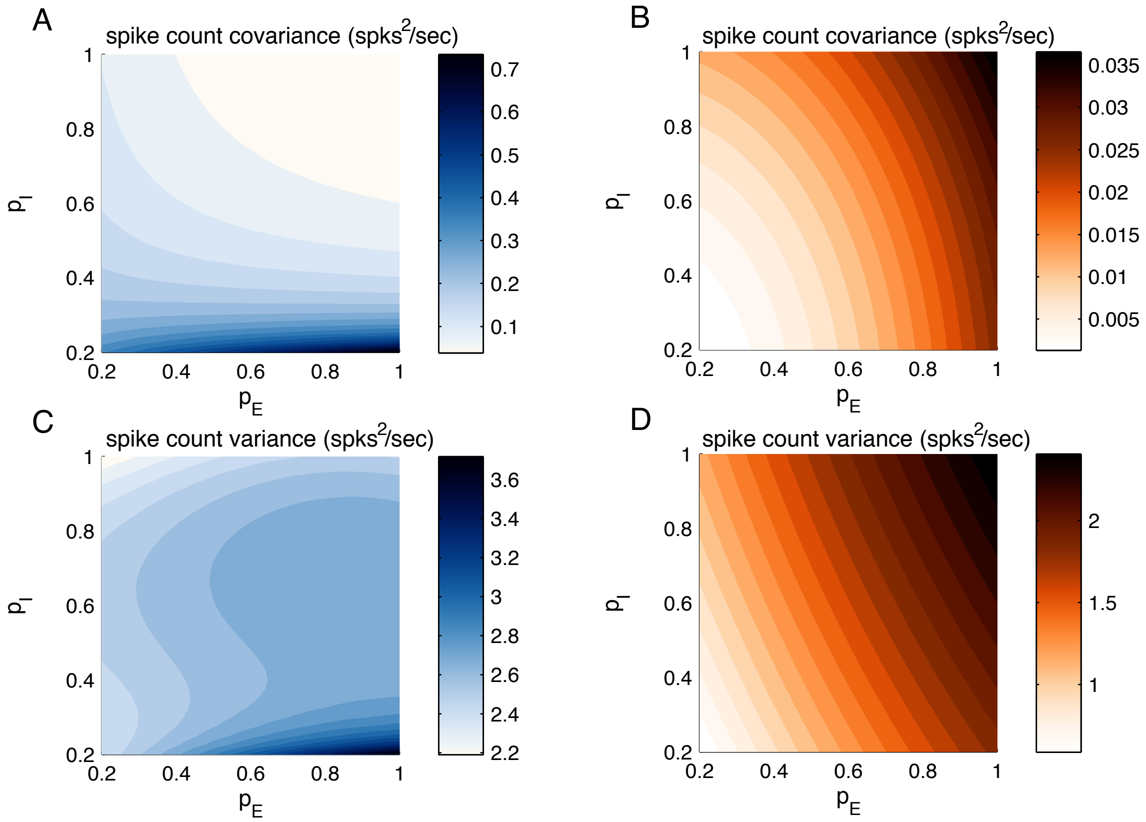


**Figure 1:** Differential changes in spike count covariance and variance due to external or recurrent changes in synaptic reliability. (**A,C**) Theoretical calculations of excitatory neuronal spike count statistics as recurrent synaptic probability of release varies. (**A**) Population averaged spike count covariance, varying excitatory ($p_E$) and inhibitory ($p_I$) probability of release separately. (**C**) Population average spike count variance, same as in A. (**B,D**) Theoretical calculations of excitatory neuronal spike count statistics as external synaptic probability of release varies. (**B**) Population averaged spike count covariance, varying excitatory ($p_E$) and inhibitory ($p_I$) probability of release separately. (**D**) Population average spike count variance, same as in B.

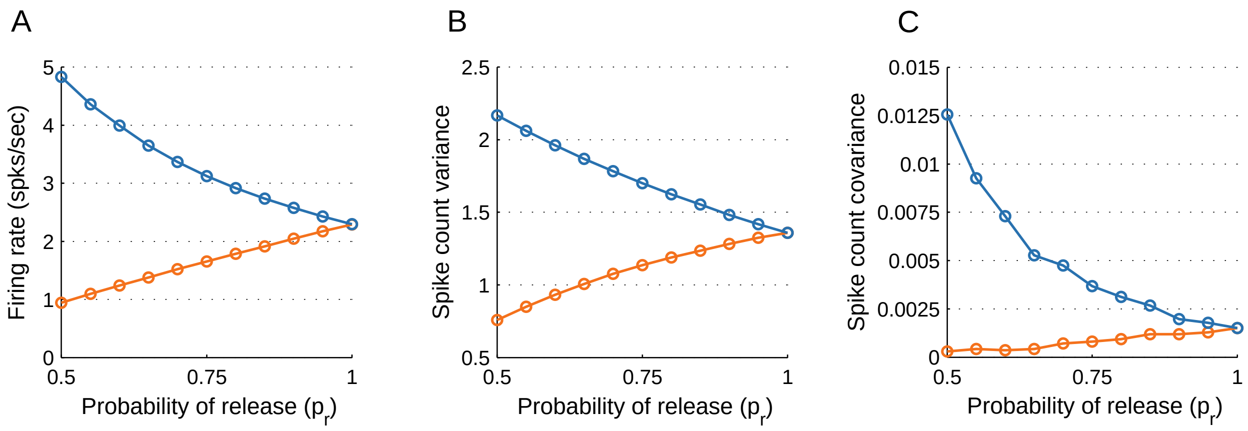


**Figure 2:** Population averaged spike count statistics of excitatory neurons in the exponential integrate-and-fire model. Simulations of the network for different values of probability of release. (**A**) Firing rate. (**B**) Spike count variance. (**C**) Spike count covariance.
